# Prot-SCL: State of the art Prediction of Protein Subcellular Localization from Primary Sequence Using Contrastive Learning

**DOI:** 10.1101/2023.09.01.555932

**Authors:** Sam Giannakoulias, John J. Ferrie, Andrew Apicello, Carter Mitchell

## Abstract

Protein subcellular localization is a critically important parameter to consider when designing expression constructs and production strategies for industry scale protein production. In this study, we present Prot-SCL an innovative self-supervised machine learning approach to predict protein subcellular localization exclusively from primary sequence. The models herein were learned from a dataset of subcellular localizations derived by exhaustively analyzing the Uniprot database. The set of localization data was rigorously curated for machine learning by employing group sampling following clustering of the protein sequences. The novel component of this approach lies in the development of a triplet neural network architecture capable of generating meaningful embeddings for classification of protein subcellular localization. We observed a robust predictive power for our classical gradient boosted machine learning models trained on these triplet embeddings in both cross validation and in generalization to the testing set. Importantly, we have made this extensive dataset of protein subcellular localizations publicly accessible, facilitating future, need-based, localization studies. Finally, we provide the relevant codebase to encourage a wider adoption and expansion of this methodology.

**GRAPHICAL ABSTRACT:** 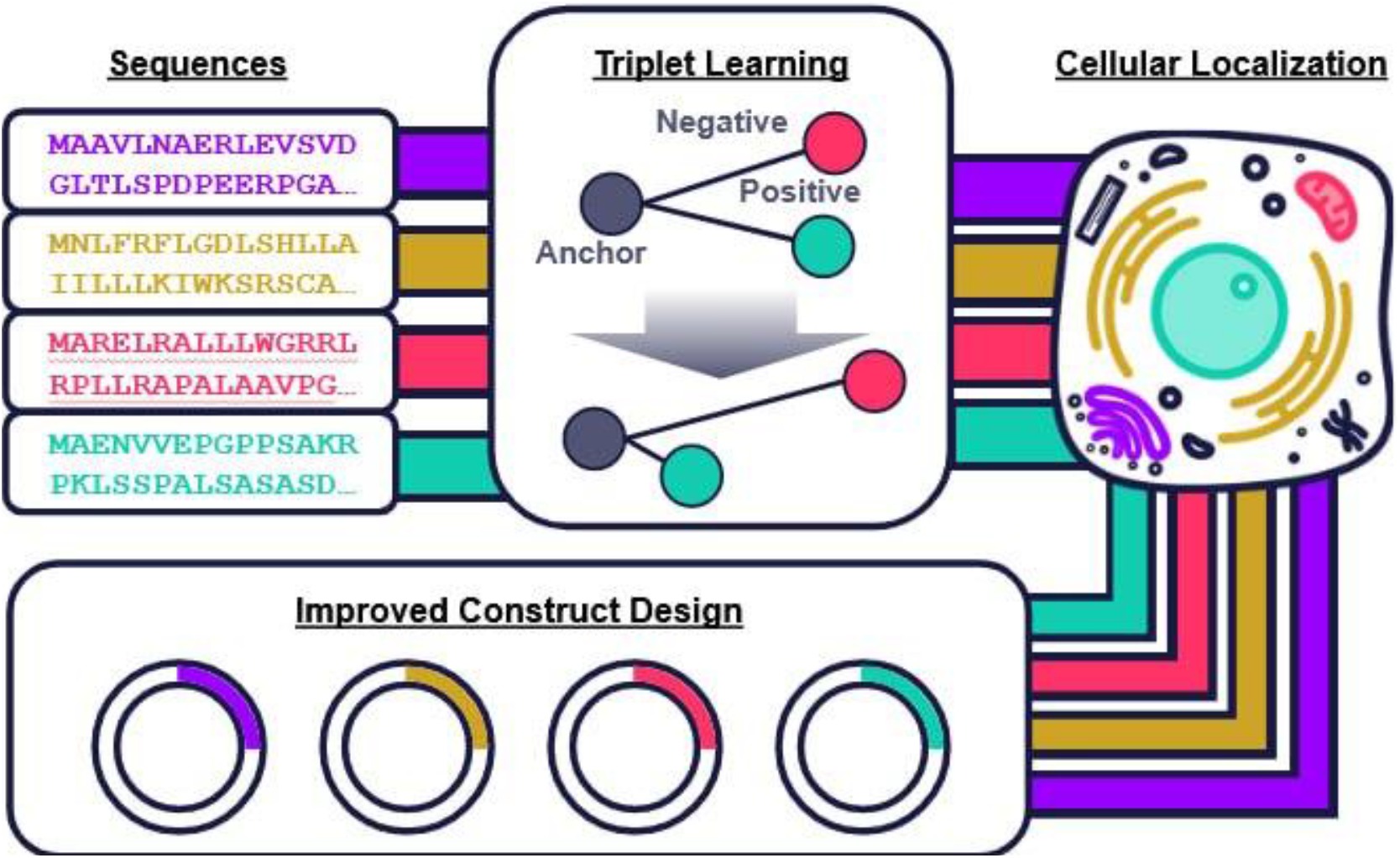

## INTRODUCTION

Protein subcellular localization constitutes a remarkably intricate cellular organization system with profound implications spanning protein functionality, disease mechanism, and biotechnology applications (1-3). The dynamic, multi-compartment, localization of proteins by intrinsic sorting signals (such as nuclear localization and export signals) and post translational modifications underscores the complexity of this phenomenon (4-6). It is worth noting that beyond these understood mechanisms, there likely exist numerous other cellular processes governing protein localization. Concordantly, the Uniprot database boasts an extensive array of annotated subcellular localizations (459 unique localizations in total), yet only a limited number of robust sorting signals are known (7,8). This holds true even for major organelles like the Endoplasmic Reticulum and Golgi Apparatus, where sorting signals remain largely opaque (9,10). Given these gaps in understanding, this article introduces a novel computational strategy to illuminate cellular organization intricacies and leverage this knowledge for advancing protein construct production.

When an abundance of well-annotated biological data is available, bioinformatic and machine learning approaches have emerged as highly effective strategies to uncover hidden patterns (11,12). Recently, in line with the advent of the transformer architecture (13), self-supervised deep learning models, namely Protein Language Models (PLMs) have become extremely powerful computational biology tools (13-15). Most PLMs, like ProtBERT (14) or ESM (16) are trained with a masked language model objective (17). That is to say, these models leverage extensive corpora of protein sequences, enabling deep learning algorithms to unveil an evolutionary-like landscape of proteins by accurately filling in masked amino acids randomly incorporated within the sequences (14,16). A very powerful aspect of PLMs is the ability to extract fixed length representations of proteins from variable length sequences. These fixed length representations, called embeddings, are able to be used as input features for machine learning models.

Notably, the DeepLoc 1.0 (18) and 2.0 (19) models were the first to use PLMs to predict protein subcellular localization. Historically, models prior to DeepLoc such as YLoc+ (20) or PROSITE (21) utilized features like sorting signals and gene ontology annotations (22). Although PLMs demonstrated a modest improvement in the ability to predict protein subcellular localization, the reported generalizability of these models to the testing set was poor. However, it was unclear whether this drop-off in predictivity was truly due to lack of generalizability or suboptimal construction of a test set which exhibited significant data drift from the training data. Therefore, this work focuses on both construction of our own dataset from the Uniprot database and the development of new PLM-based strategies for predicting these data.

## MATERIAL AND METHODS

### Dataset Construction

The dataset used for the machine learning prediction of protein subcellular localization was derived from the Uniprot database (7). Specifically, the entire reviewed Swiss-Prot subset was downloaded as xml from (https://ftp.uniprot.org/pub/databases/uniprot/current_release/knowledgebase/complete/uniprot_sprot.xml.gz) for processing. The SeqIO module from BioPython library (23) was used to parse the xml file where information about accession sequence, organism, and subcellular localization annotations were recorded. In total, we identified 569,516 unique accessions with at least one annotated subcellular localization. This included 459 unique subcellular localizations. A csv file containing every subcellular annotation with the corresponding number of accessions can be found on our GitHub at https://github.com/Sentauri-OpenSource/SubcellularLocalization/blob/main/Data/SubcellularLocalization_Counts_082020 23.csv. Additionally, a saved json file containing the subcellular localization data from the Swiss-Prot subset can be found at https://github.com/Sentauri-OpenSource/SubcellularLocalization/blob/main/Data/SubcellularLocalizations_WithAccessions_08202023.json. For the purposes of this study, we focused on the human proteome and the following subcellular localizations: Nucleus, Mitochondria, Endoplasmic Reticulum, Golgi Apparatus, Cytoplasm, Cell Membrane, and Extracellular region. For clarity, more detailed subcellular localizations, such as inner mitochondrial membrane, were associated with the previously listed more generic localizations, in this case Mitochondria. These filters left us with a considerable sized dataset of 16,632 human accessions. We have kept all other organismal and subcellular localization data present in the above-mentioned files for others with different interests and needs.

### Featurization By Protein Language Model

Once the custom dataset was generated, input sequences were featurized for future use in machine learning. The transformers and tokenizers Python libraries from HuggingFace (24) were used to embed the protein sequences. Specifically, the model weights for the ProtBERT model (14) were downloaded (upon import) for embedding. The ProtBERT model was instantiated as an instance of BertForSequenceClassification and the pretrained method was set with the value ‘Rostlab/prot_bert’ and output_hidden_states=True flag. Protein sequences were then space separated for tokenization with the BertTokenizer object which also set the pretrained method with ‘Rostlab/prot_bert’. When the tokenizer was invoked, it was done so setting padding=‘max_length’, truncation=True, and max_length=2048. This cutoff of 2048 residues was performed to reduce memory demand and was appropriate as this filter retained > 99.9% of the available data. Tokenized sequences were then fed to the model and hidden states were processed according to the following procedure: transposition of the batch size and attention layer axes, summation of features along the attention layer axis, followed by averaging over the sequence length. This process produces a PyTorch (25) tensor of shape batch size by 1024 which corresponds to the number of features per token in the ProtBERT model. All batched embeddings were then concatenated as the featurized custom dataset. The featurization script can be found at https://github.com/Sentauri-OpenSource/SubcellularLocalization/blob/main/ProtBERT_Featurization/get_protbert_embeddings.py.

### Sequence Clustering

Following featurization by ProtBERT, the custom dataset was then in a form suitable for unsupervised machine learning clustering to develop rigorous training, validation, and testing sets for supervised machine learning. The Density Based Spatial Clustering of Applications with Noise (DBSCAN) from scikit-learn (26) was chosen as the algorithm to cluster human ProtBERT embeddings. The DBSCAN model was tuned with the Optuna Python library (27) according to the following constraints. A conditional expression enforced the criterion that no individual cluster including the noise cluster contain greater than 50% of the total samples. Beyond this conditional, the eps, min_samples, and p parameters were optimized with the Bayesian optimization TPE sampler with the objective to minimize the product of the size of the noise cluster with the Davies-Bouldin score (28). cluster assignments from the minimum scoring trial of 1000 trials were used for clustering sampling. Training, validation, and testing splits were constructed according to an approximate 80-10-10 distribution. Validation and testing sets were populated with randomly selected clusters, taking all of their samples. This ensured that there is no imminent redundancy of samples between machine learning sets in the domain of the features used for machine learning. The final criterion was that the noise cluster was ensured to be contained in the training set to provide these important samples to the learning process. Clustering and sampling scripts can be found at https://github.com/Sentauri-OpenSource/SubcellularLocalization/blob/main/ProtBERT_Clustering/cluster_sequences_dbscan.py.

### Triplet Generation

The machine learning sets were then used to create triplet samples. Triplets were defined such that every sample in the data would be used as an anchor. Positive selections were enforced such that identical subcellular localization profiles were observed. For instance, if a sample is found in the Endoplasmic Reticulum and also the Cytoplasm, and another sample is only found in the Cytoplasm, these two samples were not considered to be positive examples as they did not share identical localization profiles. Negative examples were enforced such that they share exactly zero subcellular localizations in common with the anchor sample. The number of triplets generated per anchor was non-fixed as determined by the number of subcellular localizations of the anchor sample. Random sampling from the entirety of the per-anchor negative pool was performed until a sample from every anchor non-localizing compartment was selected making sure to not duplicate any such negative example subcellular localizations. Finally, the number of unique positive examples per anchor triplet set was dictated by the number of negative examples and the existence of identical positive examples. If there were sufficiently many identical positive examples to produce a completely unique set of anchor-positive-negative triplets, this was done. If there were not, positives were selected with replacement. Triplet generation scripts can be found at https://github.com/Sentauri-OpenSource/SubcellularLocalization/blob/main/Offline_Triplet_Generation/make_triplets_offline.py.

### Triplet Representation Learning

A custom PyTorch triplet neural network script was written to learn representations of the protein sequences which were more representative of protein subcellular localization than simply ProtBERT embeddings. Again, the Optuna Python library was used to hypertune the triplet neural network. In particular, the number of linear layers, the decay rate (a custom hyperparameter used to determine how quickly the number of layer units will decrease), the final embedding dimension, activation function, dropout weight, initial learning rate, learning rate scheduler gamma value, and batch size were tunable hyperparameters. Given the already large sized dataset and the significant increase in size from triplet generation, we chose to fix the number of epochs at 2 with early stopping available. Early stopping was constructed with a patience value of 2 where validation was performed every 10^th^ of an epoch. The triplet neural network was tuned to minimize the triplet loss of the validation set triplets using the TPE sampler as before. Model training scripts can be found at https://github.com/Sentauri-OpenSource/SubcellularLocalization/blob/main/Triplet_Representation_Learning/train_subcellular_localization_triplet_network.py.

### Supervised Learning

The optimal triplet neural network model was then used to extract embeddings for use in unsupervised and supervised machine learning. In this way, we have effectively gone from variable length protein sequences to a fixed length ProtBERT representation and then finally to a new, smaller, fixed length subcellular localization informed embedding. These informed embeddings were then put into an unsupervised dimensionality reduction routine with the UMAP Python library (29). The embeddings were projected into 2 dimensions and colored according to their subcellular localization with the matplotlib Python library (30). Additionally, embeddings from both the training (training and validation sets combined) and testing sets were assessed by the Davies-Bouldin score. Beyond these unsupervised analyses, we trained gradient boosted tree models for each subcellular localization task, given the tabular nature of the embedding data (final learning set concatenated triplet embeddings with ProtBERT embeddings). The CatBoost Python library (31) was chosen as the flavor of gradient boosted tree model and, again, the Optuna Python library was employed to hypertune these models. For each of the subcellular localization dependent variables, we hypertuned the following hyperparameters for 100 trials with the TPE sampler: grow_policy, iterations, learning_rate, depth, l2_leaf_reg, bagging_temperature, random_strength, and scale_pos_weight. The Optuna objective here was to maximize the F1 score, a metric which reports on both the precision and recall of the minority class and is suitable in situations of class imbalance. The F1 value here represents the true F1 from the concatenation of all predictions from five-fold StratifiedGroupKFold cross validation. The CatBoost training scripts can be found at https://github.com/Sentauri-OpenSource/SubcellularLocalization/tree/main/Supervised_Training.

## RESULTS

### Protein Subcellular Localization Dataset

To develop robust predictors of protein subcellular localization, we constructed a custom dataset from the experimentally validated Swiss-Prot subset of Uniprot. The Swiss-Prot set was processed according to organism and presence of at least one subcellular location. Raw exploratory data analysis metrics are presented in Table 1. We used the human only dataset with major compartment localizations as the dataset for machine learning. These sequences were curated into training, validation, and testing sets based on sampling from cluster assignments from the DBSCAN algorithm. DBSCAN was chosen over alternative clustering algorithms such as KMeans or Gaussian Mixture Models as to not assert the number of clusters the data should be labeled to. Additionally, the noise cluster from DBSCAN represents a rich set of unique points which can be utilized for learning in the training set of our supervised models. The breakdown of dependent variable labels in each machine learning dataset can be seen in Table 2. It should be noted that the sum of these columns will not exactly reproduce the 80-10-10 split in terms of number of accessions, as many sequences contain multiple subcellular localizations.

**Table 1.** Table exhibiting the number of unique organisms, subcellular localizations, and sequences in the unmodified Swiss-Prot set, the subset of Swiss-Prot with at least one subcellular localization, and in the human only and at least one subcellular localization set. The seven major compartments which represent the dataset for this study include the Nucleus, Mitochondria, Endoplasmic Reticulum, Golgi Apparatus, Cytoplasm, Cell Membrane, and Extracellular region.

TABLE AND FIGURES LEGENDS
| Dataset | Num. Organisms | Num. Localizations | Num. Accessions |
| --- | --- | --- | --- |
| Swiss-Prot | 14439 | > 459 | 569,516 |
| Swiss-Prot<br>with Localization | 12,589 | 459 | 354,887 |
| Swiss-Prot Human<br>with Localization | 1 | 242 | 16,632 |
| Swiss-Prot Human with<br>Localization to Major<br>Compartments | 1 | 7 | 16,632 |

**Table 2.** Table displaying the number of positive examples for each dependent variable label across all machine learning sets.

| Label | Training Set (#) | Validation Set (#) | Testing Set (#) |
| --- | --- | --- | --- |
| Nucleus | 4,348 | 422 | 555 |
| Mitochondria | 1,120 | 78 | 90 |
| Endoplasmic<br>Reticulum | 1,036 | 98 | 139 |
| Golgi Apparatus | 734 | 79 | 110 |
| Cytoplasm | 4,255 | 482 | 623 |
| Cell Membrane | 3,026 | 371 | 308 |
| Extracellular Region | 1,526 | 338 | 189 |

### Prediction of Subcellular Localization with ProtBERT Embeddings

With curated data sets in hand, we performed supervised machine learning directly on the ProtBERT embeddings. This experiment represents the easiest implementable solution and provides a baseline control for more sophisticated methods. We observed that for subcellular localizations with large support (number of examples), such as Nuclear localization, ProtBERT embeddings were quite effective machine learning features (accurate and low overfitting). However, for lower support localizations such as the Golgi Apparatus, these supervised machine learning models exhibited significantly poorer performance especially to the testing set (less accurate and more overfitting).

Historically, literature models have sought to improve the quality of their models by incorporating sorting signal information to their machine learning. We investigated the sorting signal dataset used by DeepLoc and uncovered many issues. Firstly, the number of sequences which contained annotated sorting signals was substantially smaller than those with subcellular localizations (<10%). Additionally, we observed there were a large number of inconsistencies between the DeepLoc Swiss-Prot dependent variable annotations and the presence of sorting signals and vice versa. When we then simply compared the DeepLoc subcellular localizations for the same accessions as our processed localizations dataset, we observed even more differences. It is unclear the reason for these differences beyond the fact that we used a completely up to date Swiss-Prot dataset. This phenomenon demonstrates the imperative need for bioinformaticians and data scientists who train models built upon annotation data to be very heavily monitored and updated with the latest burgeoning research. With that being said, we took inspiration from the idea of using sorting signals and sought to develop a contrastive learning-based learning approach which could mitigate the need for annotated sorting signals.

### Triplet Learning

Contrastive learning is a well-known self-supervised machine learning technique which makes use of sample relationships to develop robust representations. The most common contrastive learning methods are Siamese (considers two samples at once), or Triplet (considers three samples at once) Neural Networks. Here we trained a Triplet neural network based on subcellular localization using only sequence level ProtBERT embeddings as feature inputs. The goal of the Triplet network is to show the model many examples of similar and dissimilar subcellular localizations, so that patterns within ProtBERT embeddings can be identified as corresponding to subcellular localizations. This is especially important for low support and low annotation localizations like the Golgi Apparatus. Figure 1. displays two representative examples of our Triplet embeddings. These data clearly show that the Nucleus (top row) and the Extracellular Region (bottom row) sample embeddings occupy unique positions on the Triplet manifold (low dimensional representation of high dimensional data). Additionally, we observe that the tuned Triplet model testing set embeddings follow a very similar distribution to the training samples (right column vs left column). This is exemplified by the small and consistent values for the Davies-Bouldin score across sets and dependent variables. All dependent variable Triplet embeddings can be found in the Supplementary Figures S1-S7.

**Figure 1.**
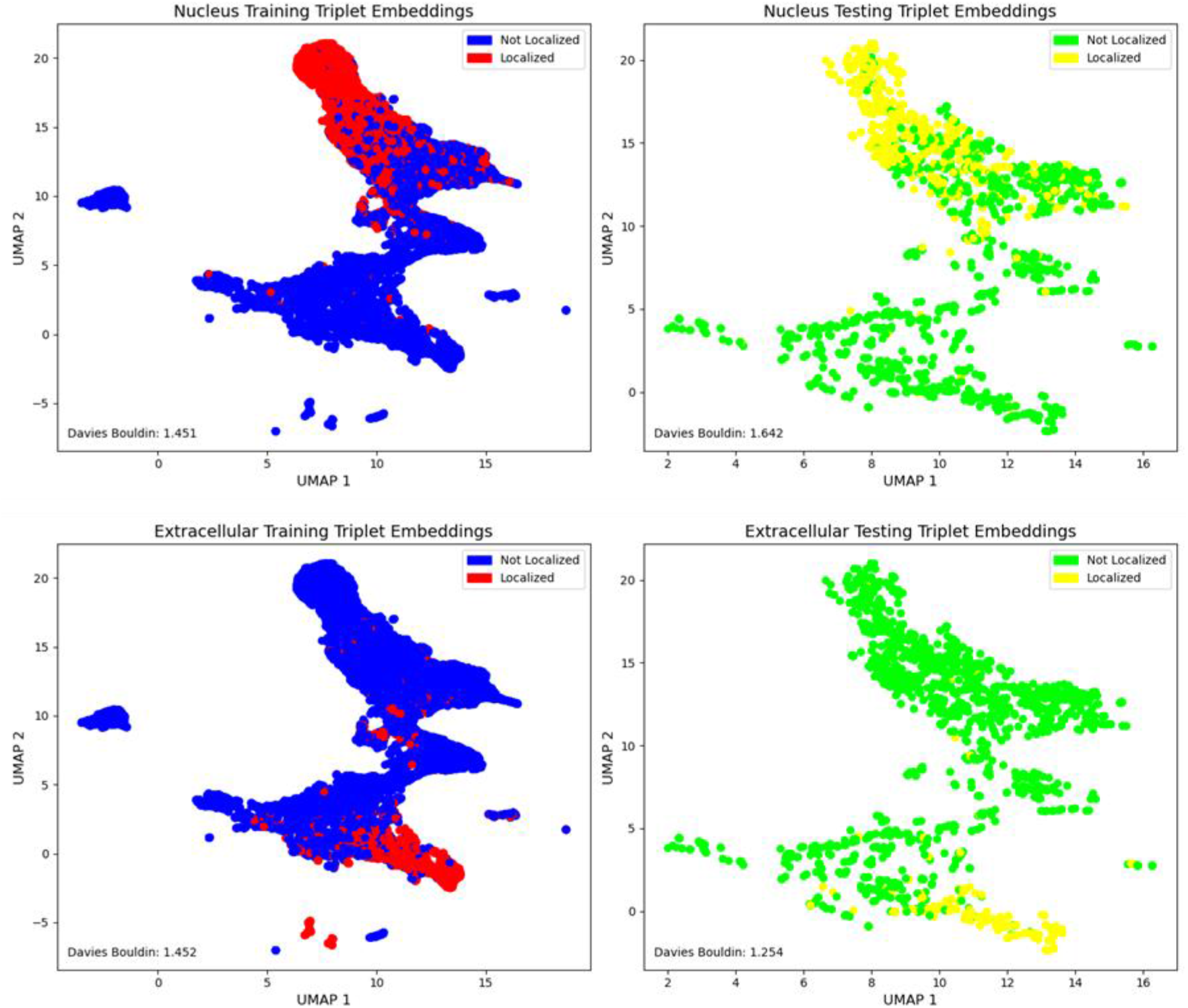
UMAP Triplet model embeddings for training (left column) and testing (right column) sets. Top row corresponds to the Nuclear subcellular localization where points in blue are non-localized and those in red are localized to the Nucleus. The same coloring is true for the bottom left representing the localization or non-localization of sequences to the Extracellular region. For testing set embeddings, green colored points represent non-localization whereas yellow points represent localization.

### Prediction of Subcellular Localization with Triplet Embeddings

Triplet model embeddings were used to train supervised machine learning models for every subcellular localization. Hypertuning parallel coordinate plots for these models can be found in the Supplementary at Figures S8-S14. Supervised classifiers were benchmarked with Receiver Operating Characteristic Curves (ROC), classification histograms, and classification reports (Figures 2 and 3, Table 3). All dependent variable subcellular localization plots can be found in the Supplementary at Figures S15-S28. We observed that similarly to the original ProtBERT supervised models, that larger support localizations performed better in the Triplet embedding supervised learning compared to lower support localizations. However, although there is still much room for improvement in predicting subcellular localizations like the Golgi Apparatus, we observed significant improvements in testing set F1 generalizability using our Triplet embeddings over purely ProtBERT embeddings. Supplementary Table S1 shows the cross validation and testing set classification reports for the ProtBERT and Triplet embedding supervised models. In the best case, the Golgi Apparatus, the use of Triplet embeddings improved the testing set F1 score by over 71%.

**Table 3.**
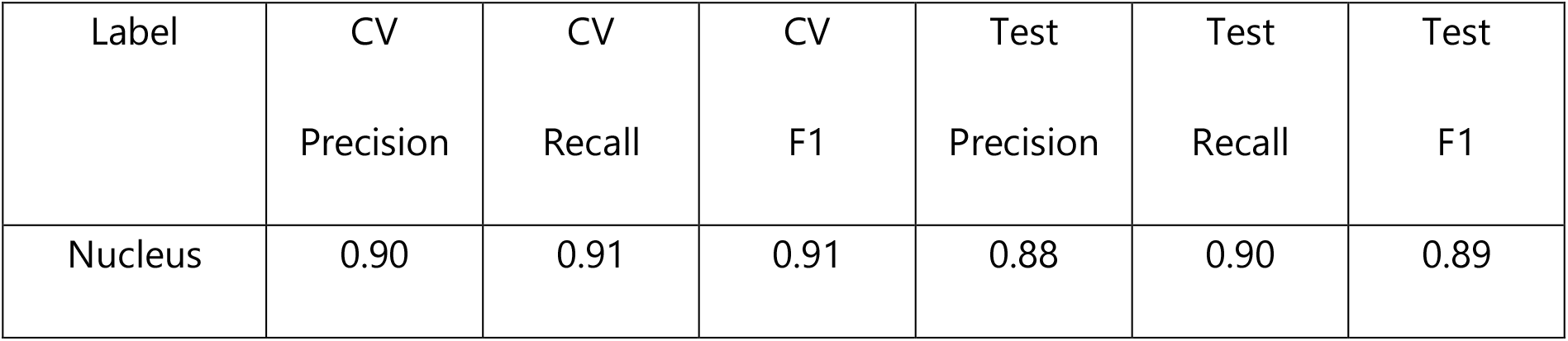
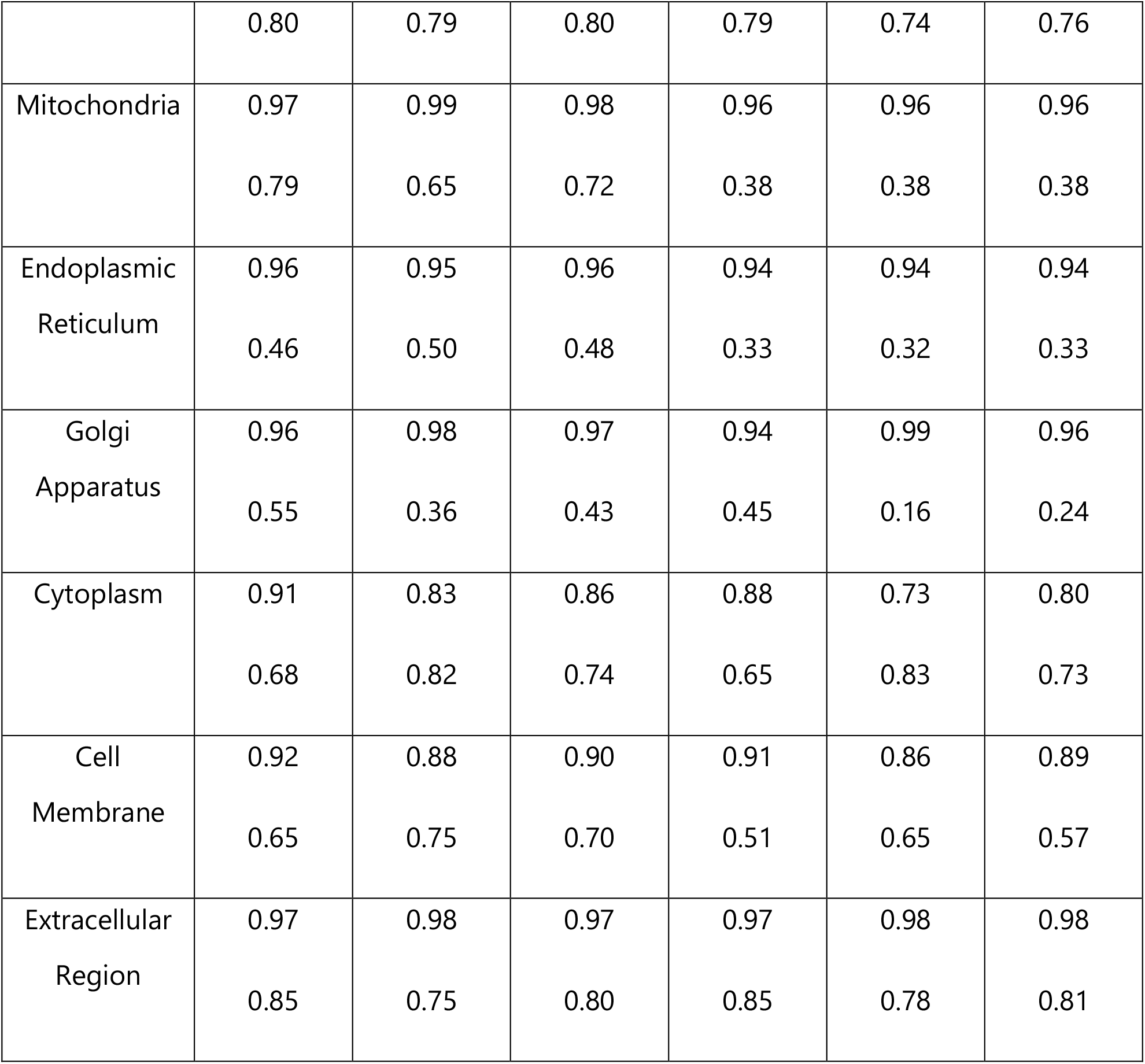
Classification reports for all supervised triplet models. Top value represents cross validation score whereas bottom value represents testing set score.

**Figure 2.**
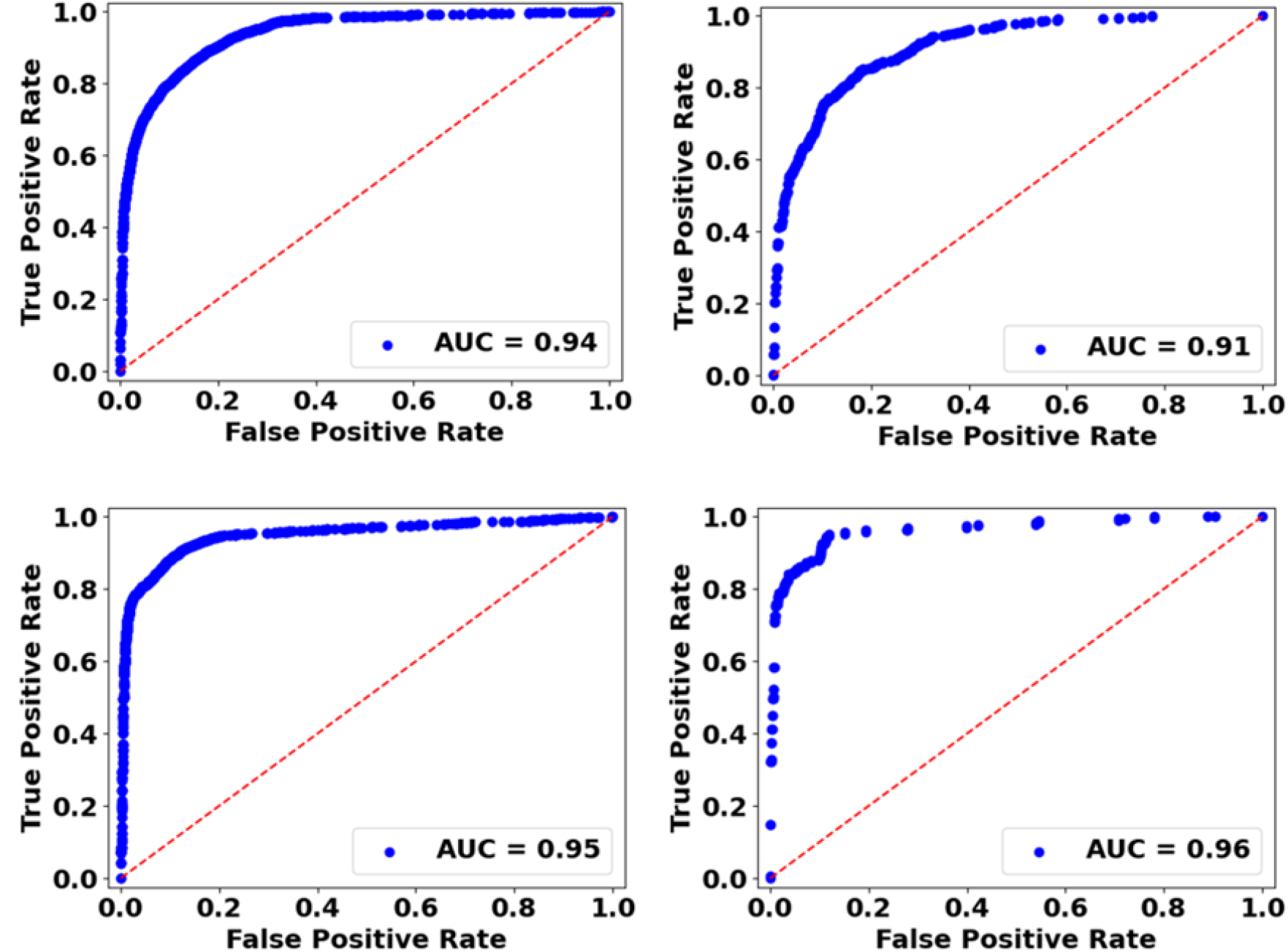
Receiver operating characteristic curves for nuclear localization (top row) and extracellular localization (bottom row). The left column displays cross validation benchmarking where the right column displays testing set benchmarking.

**Figure 3.**
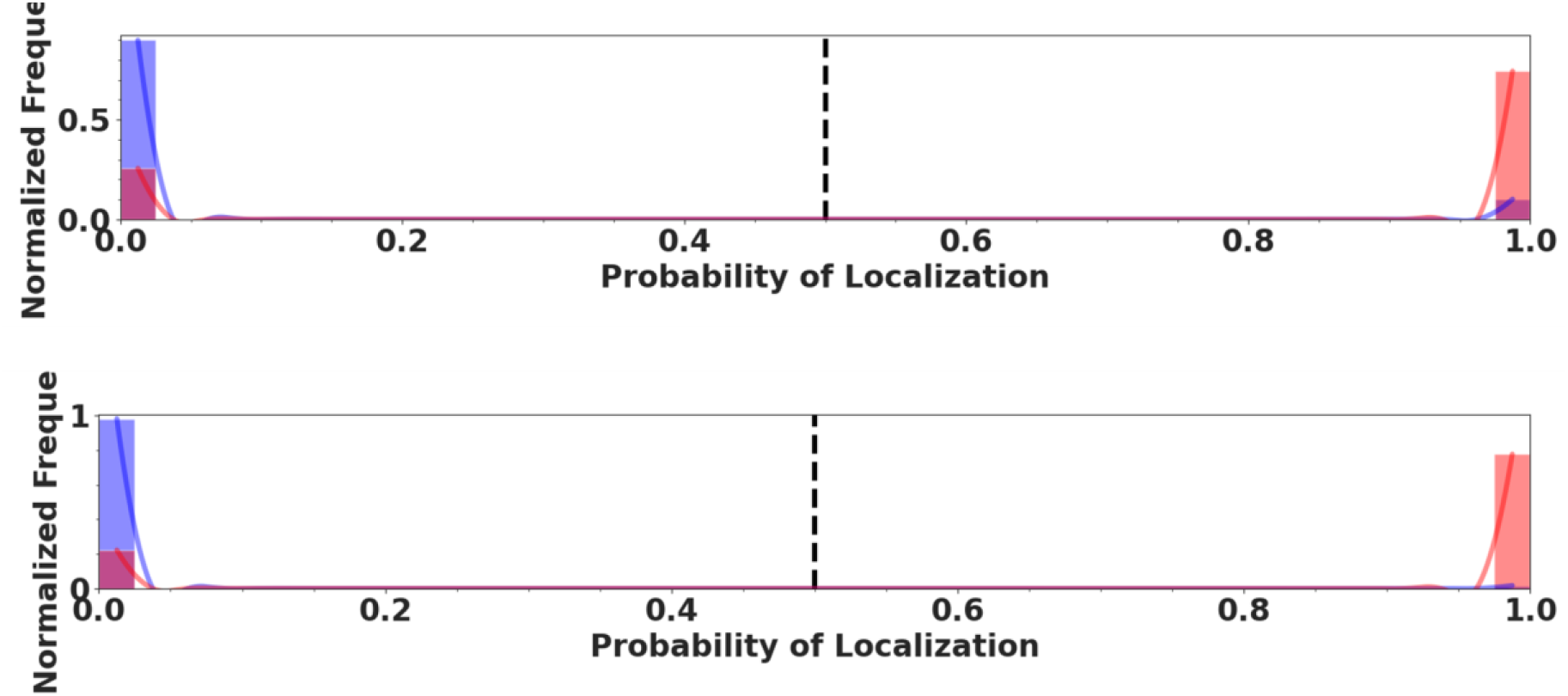
Classification histograms for nuclear localization (top row) and extracellular localization (bottom row). Red columns represent true positives, while blue bars represent true negatives. The dashed line separates the classes at a probability of 0.5.

## DISCUSSION

Here, we introduced an innovative approach for predicting protein subcellular localization exclusively from primary sequence data. By leveraging the power of self-supervised machine learning and protein language models, we developed a robust strategy to tackle the complex problem of protein localization. This work contributes to a computational foundation that can aid in academic and industrial protein production.

To build practically useful models beyond prior literature methods, this work focused on addressing the complexity of protein subcellular localization without the need for explicitly annotated sorting signals. We identified that the dataset DeepLoc 2.0 trained on utilized a very small set of annotated sorting signals as a feature engineering step ahead of the multi-layer perceptron that performs the classification. We observed that annotations between the sorting signals dataset and the greater Swiss-Prot dataset contained inconsistencies. Additionally, the labeled dataset was a significantly smaller subset than what can be found in Swiss-Prot. Furthermore, many accessions present in the DeepLoc 2.0 dataset had inconsistent labels with the Swiss-Prot dataset as of our downloading of the database in August of 2023. For these reasons we developed our own custom database for machine learning protein subcellular localization.

The custom Swiss-Prot dataset allowed us to study the ability for PLMs to predict subcellular localization. We observed that ProtBERT embeddings were able to accurately predict subcellular localizations with large amounts of available annotated data. However, localizations with lower numbers of annotations performed more poorly. We hypothesized that Triplet Neural Network architecture could be used to learn even more informative features for classification than ProtBERT embeddings by teaching a model which types of sequences were more or less similar. By doing so with a Triplet Network, we were able to utilize significantly more data on less annotated localizations than if explicit sorting signals were required. The incorporation of contrastive learning principles through our Triplet Neural Network allowed us to enhance the representation learning process and overcome limitations associated with previous models. The demonstrated effectiveness of our approach during the training gradient boosted tree models on the derived embeddings resulting in improvements in the testing set F1 scores of all but one localization.

Our work not only extends the boundaries of protein subcellular localization prediction but also has broader implications for the intersection of biology and machine learning. We have provided detailed documentation of our methodology, dataset, and codebase to facilitate further research and adoption of our approach. By making our curated dataset and source code publicly accessible, we aim to encourage collaboration, validation, and extension of our findings by the broader scientific community.

## Supporting information

Electronic Supplementary Information

## DATA AVAILABILITY

The data and all scripts required to reproduce or expand upon the work described herein can be found at https://github.com/Sentauri-OpenSource/SubcellularLocalization/tree/main.

## SUPPLEMENTARY DATA

Supplementary Data are available at NAR online.

## ACKNOWLEDGEMENTS

N/A

