## Supplementary material for "Prot-SCL: State of the art Prediction of Protein Subcellular Localization from Primary Sequence Using Contrastive Learning": Electronic Supplementary Information

**Table of Contents:**

1. **Software.......................................................................................... S2**
2. **Triplet Learning.............................................................................. S2**
3. **Supervised Learning........................................................................ S9**

### Software

The software used for this work is from one main conda environment. This environment can be installed with instructions through the following link:

https://github.com/Sentauri-OpenSource/SubcellularLocalization/tree/main/Anaconda.

### Triplet Learning

Each sample in the training, validation, and testing sets were fed to the Triplet network to extract their embeddings. These embeddings were reduced to two dimensions with UMAP and colored according to their label. These data are show in Figures S1-S7 below.

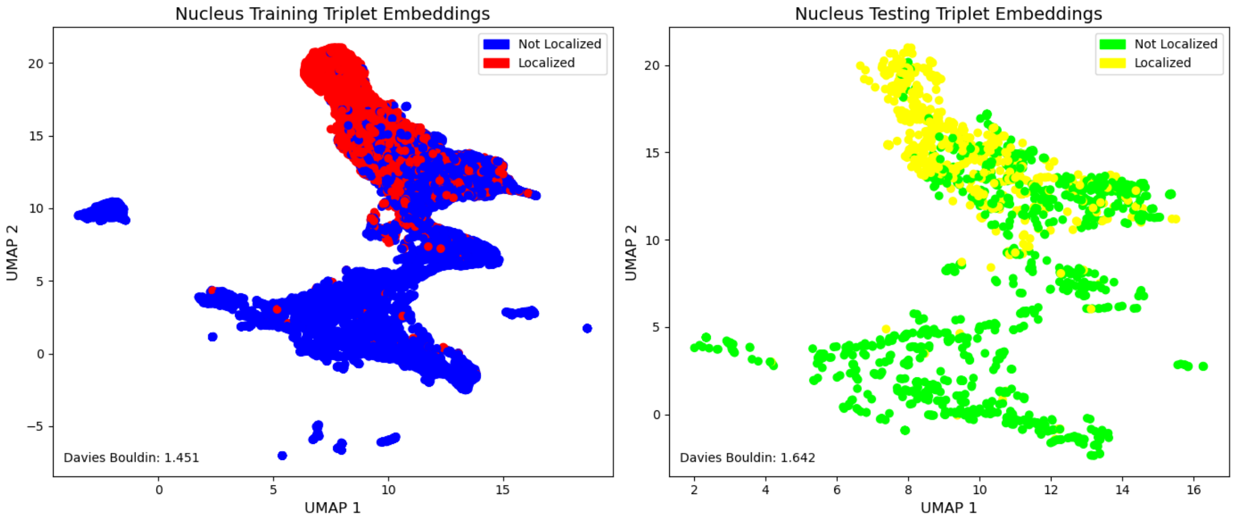


**Figure S1.** Plotted triplet embeddings colored for nuclear localization (training set on left, testing set on right).

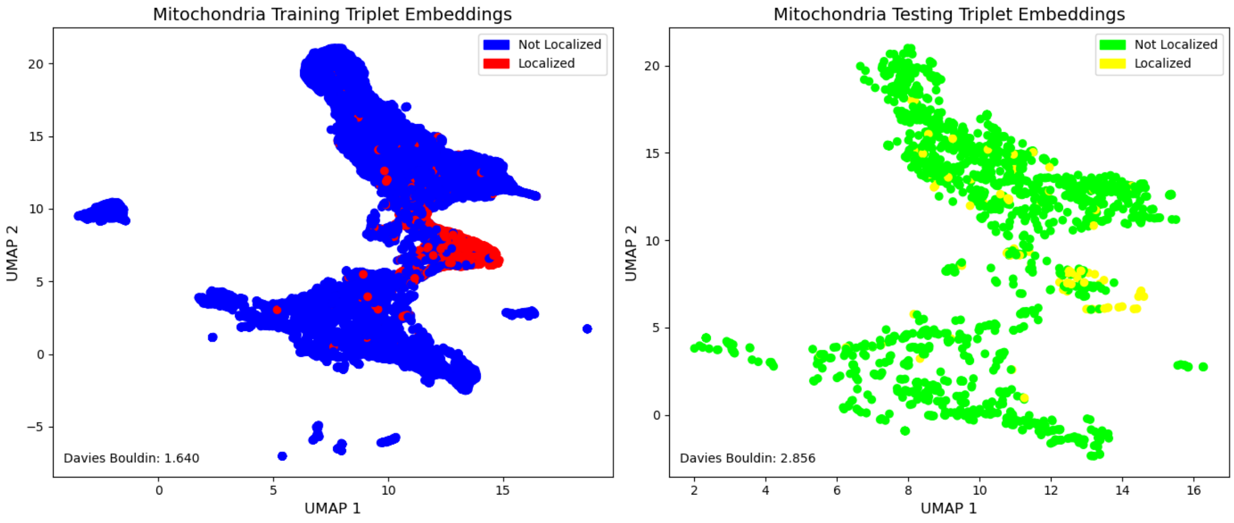


**Figure S2.** Plotted triplet embeddings colored for mitochondria localization (training set on left, testing set on right).

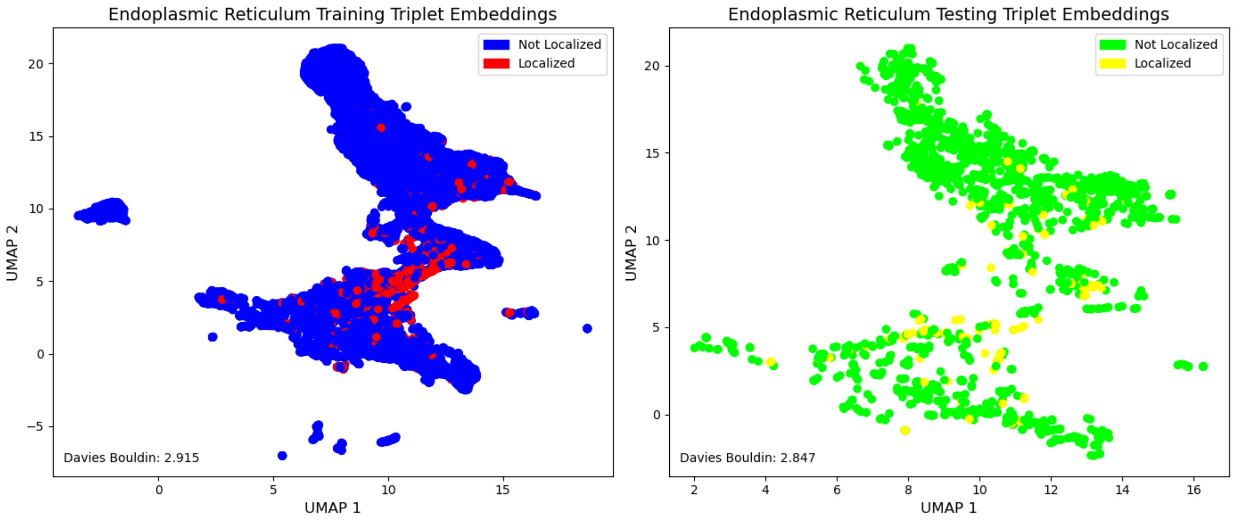


**Figure S3.** Plotted triplet embeddings colored for endoplasmic reticulum localization (training set on left, testing set on right).

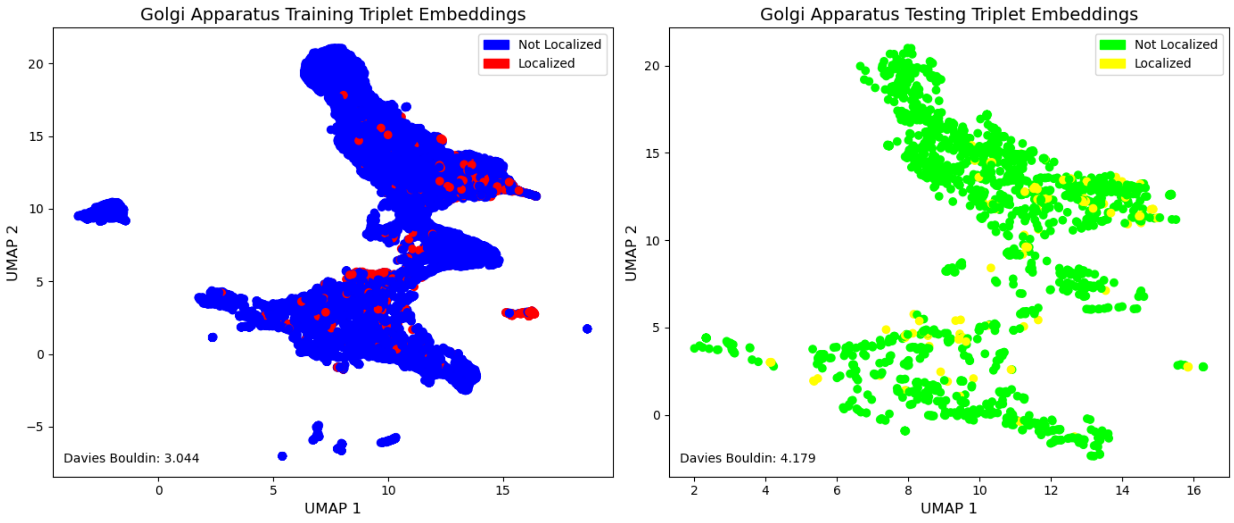


**Figure S4.** Plotted triplet embeddings colored for golgi apparatus localization (training set on left, testing set on right).

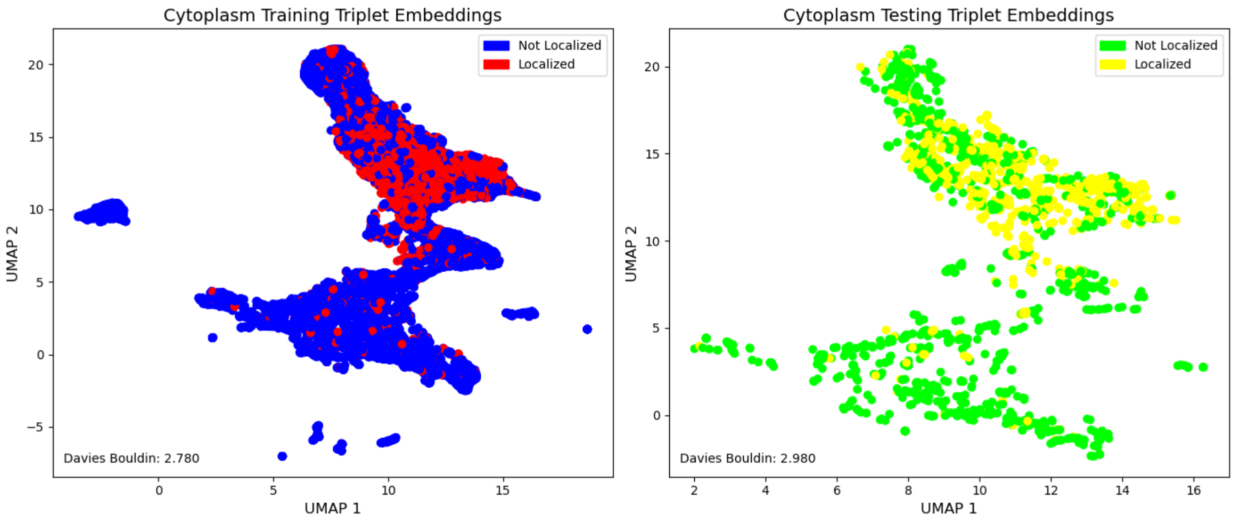


**Figure S5.** Plotted triplet embeddings colored for cytoplasmic localization (training set on left, testing set on right).

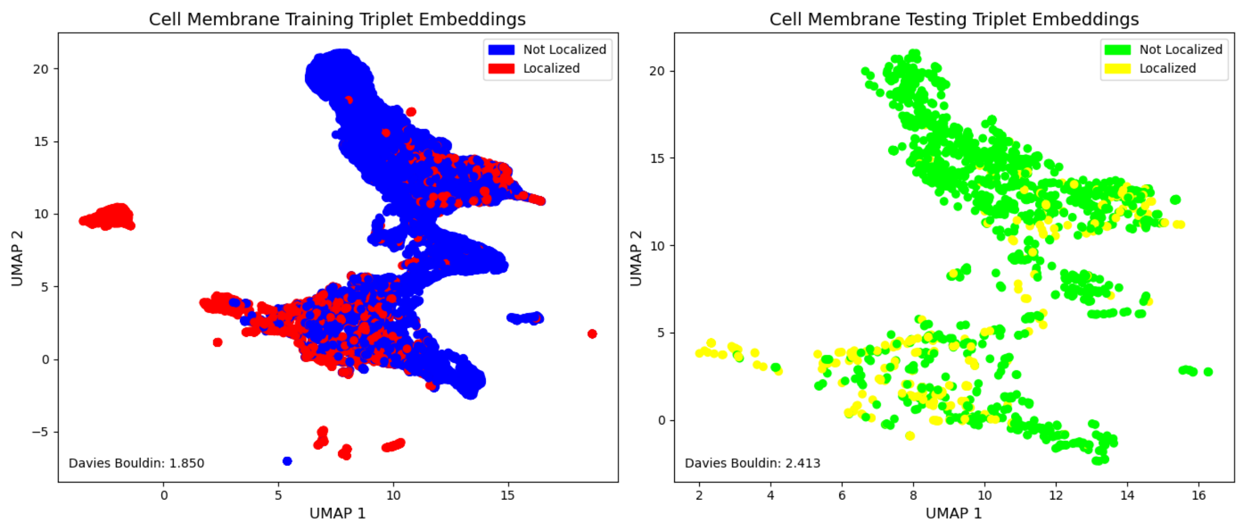


**Figure S6.** Plotted triplet embeddings colored for cell membrane localization (training set on left, testing set on right).

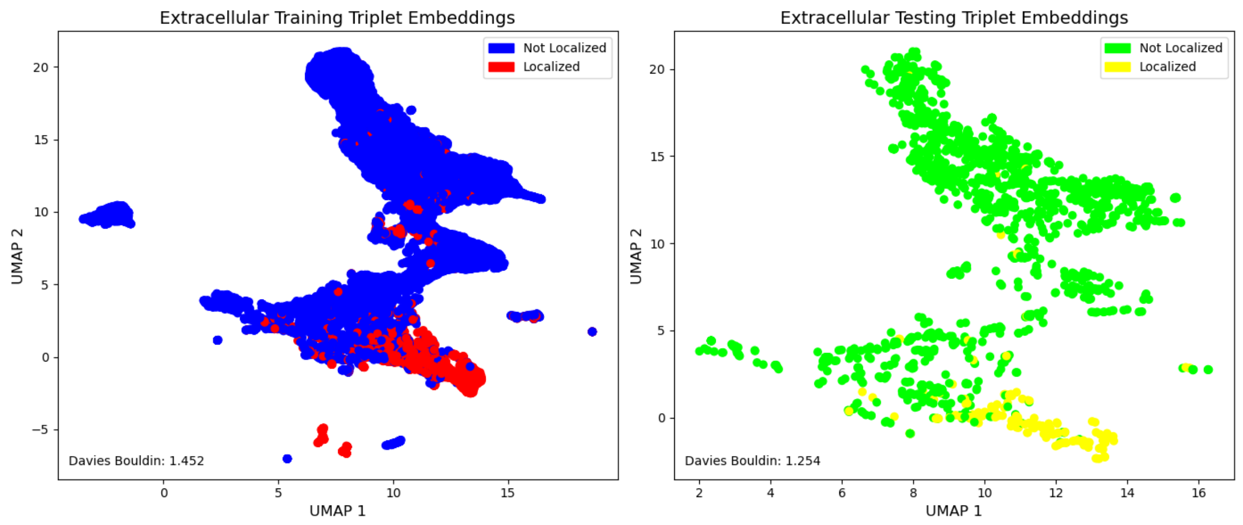


**Figure S7.** Plotted triplet embeddings colored for extracellular localization (training set on left, testing set on right).

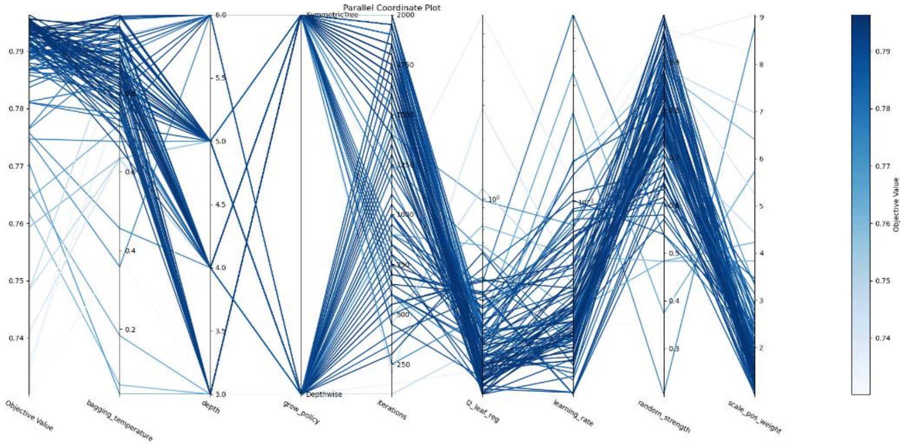


**Figure S8.** Parallel coordinate plot for nuclear localization hypertuning.

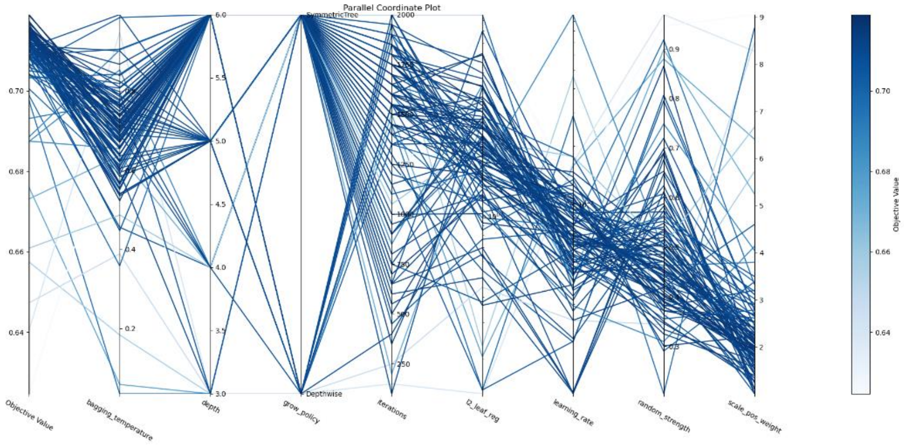


**Figure S9.** Parallel coordinate plot for mitochondria localization hypertuning.

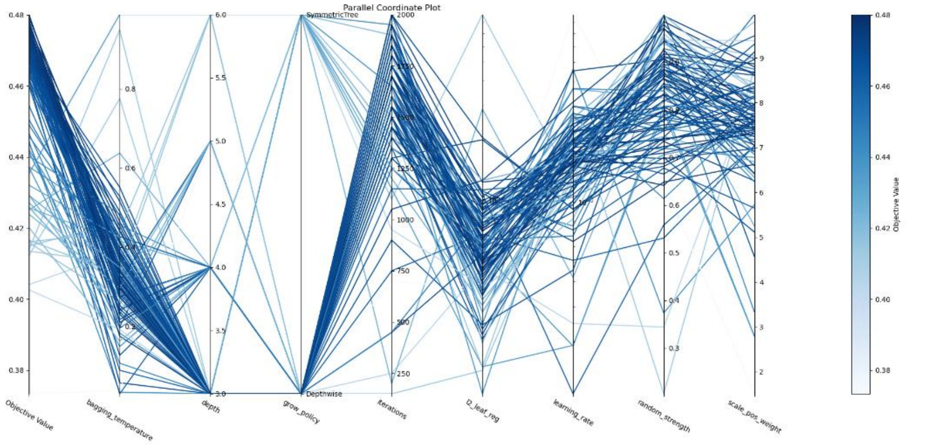

**Figure S10.** Parallel coordinate plot for endoplasmic reticulum localization hypertuning.

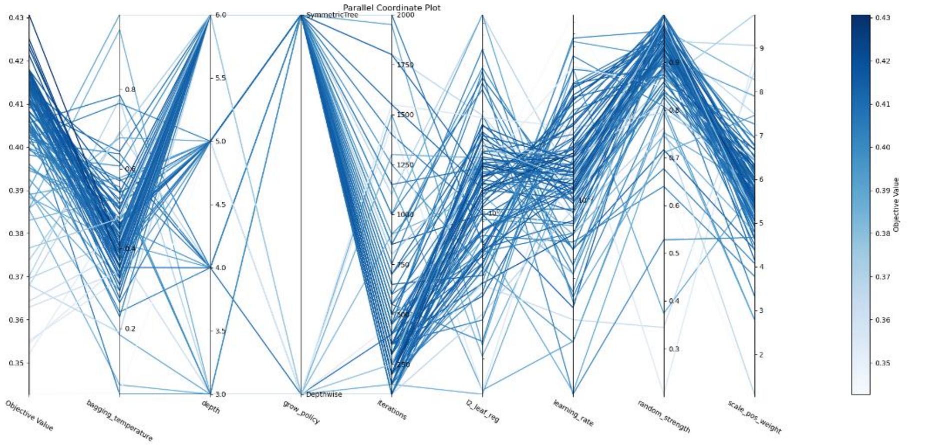

**Figure S11.** Parallel coordinate plot for golgi apparatus localization hypertuning.

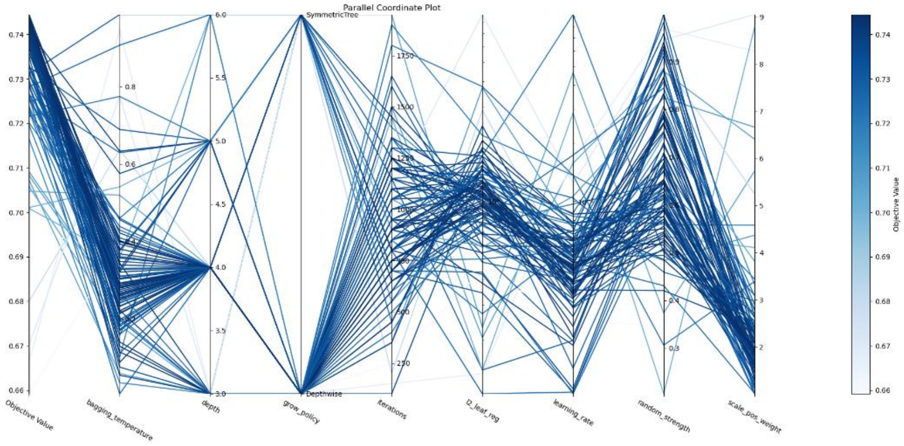


**Figure S12.** Parallel coordinate plot for cytoplasm localization hypertuning.

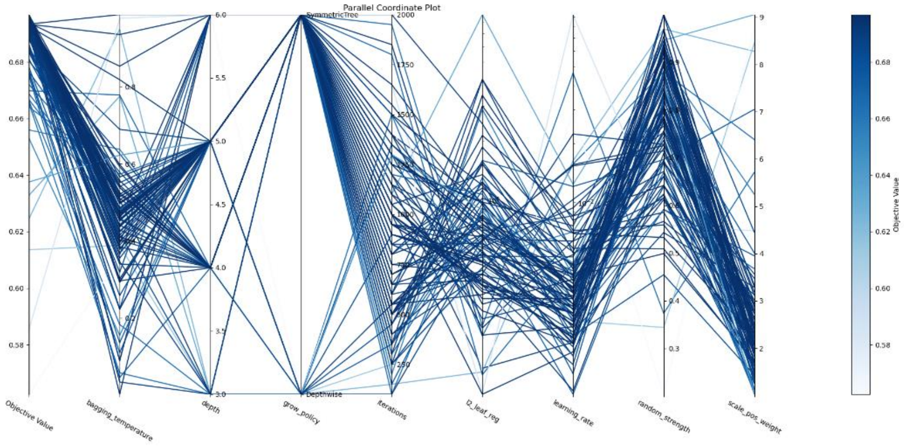


**Figure S13.** Parallel coordinate plot for cell membrane localization hypertuning.

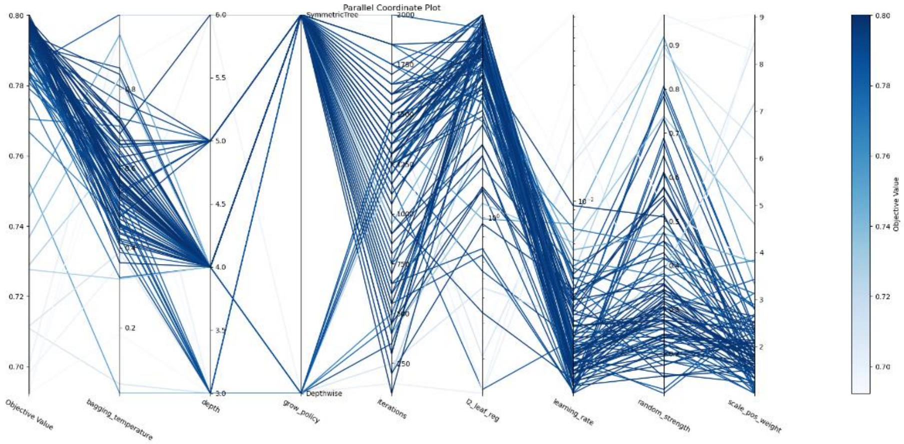


**Figure S14.** Parallel coordinate plot for extracellular localization hypertuning.

### Supervised Learning

The following section displays the full benchmarking of the supervised triplets models. These benchmarkings include receiver operating characteristic curves and classification histograms (S15-S28). Finally, Table S1 displays the % change in testing set F1 from ProtBERT embedding only models to our Triplet learning models.

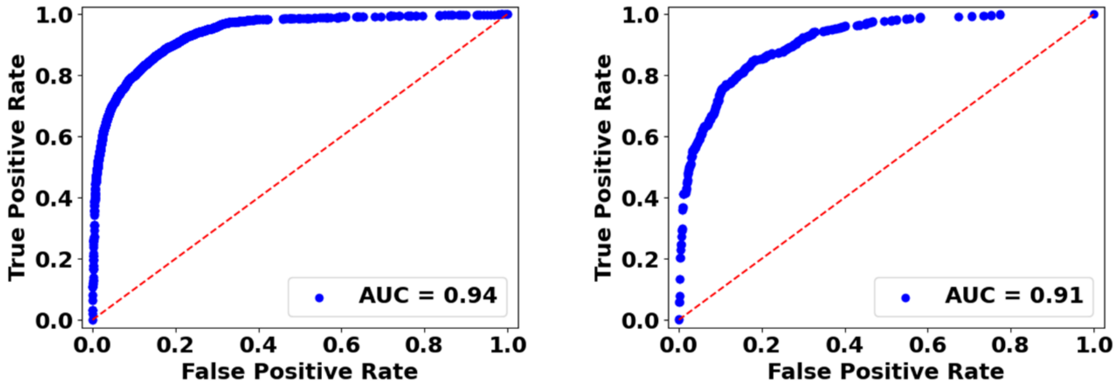


**Figure S15.** Receiver operating characteristic curve for nuclear localization (validation on left, testing on right).

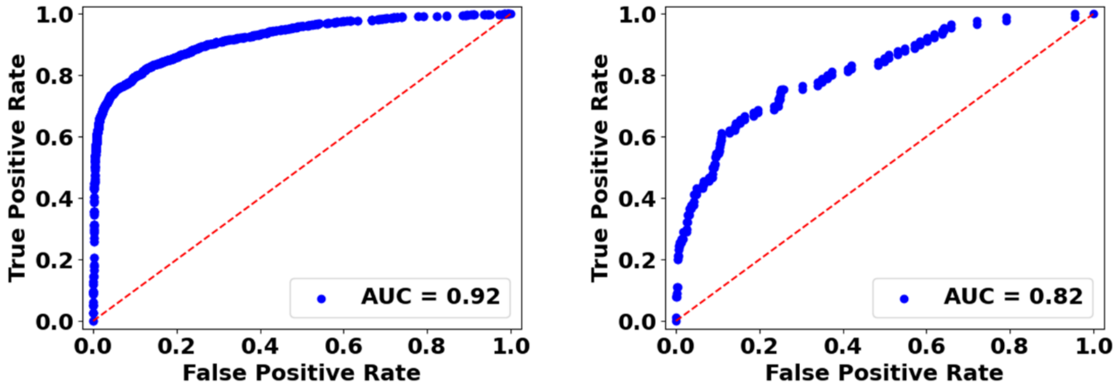


**Figure S16.** Receiver operating characteristic curve for mitochondria localization (validation on left, testing on right).

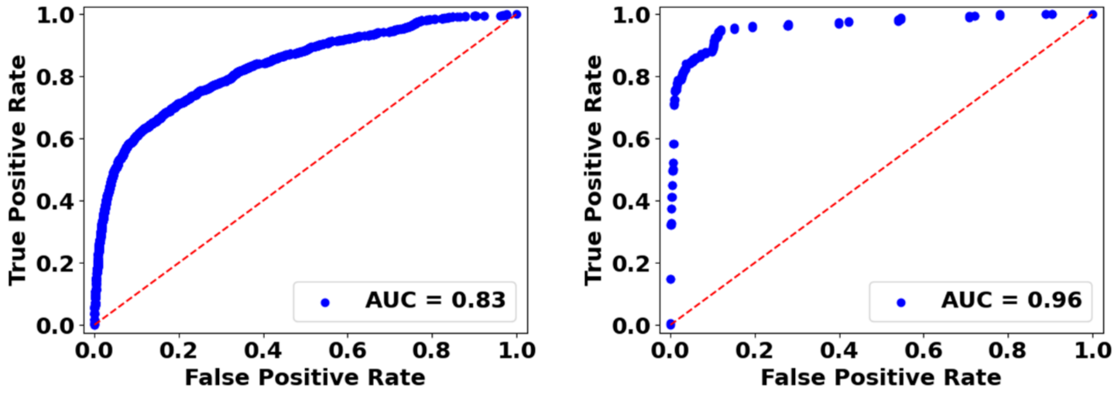


**Figure S17.** Receiver operating characteristic curve for endoplasmic reticulum localization (validation on left, testing on right).

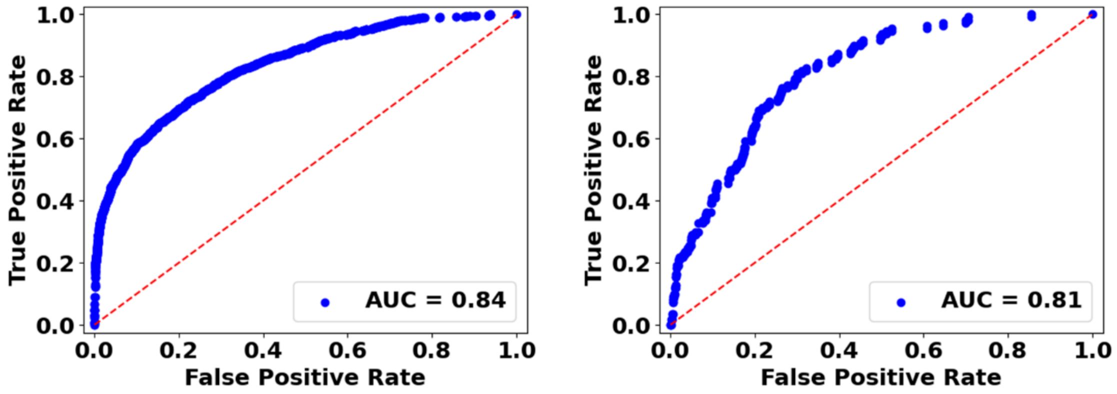


**Figure S18.** Receiver operating characteristic curve for golgi apparatus localization (validation on left, testing on right).

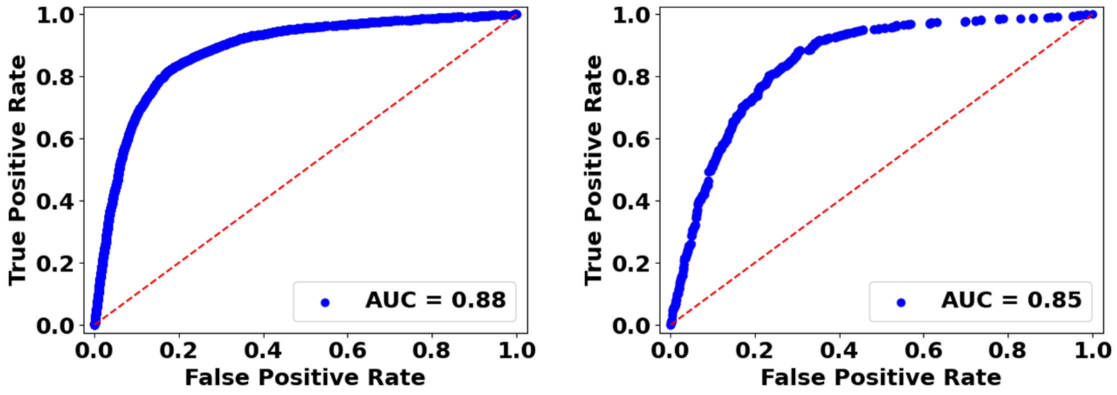


**Figure S19.** Receiver operating characteristic curve for cytoplasm localization (validation on left, testing on right).

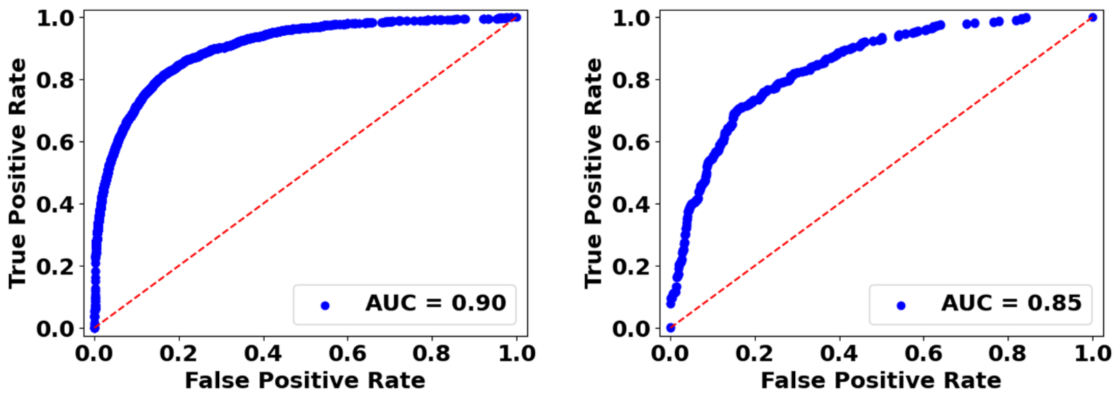


**Figure S20.** Receiver operating characteristic curve for cell membrane localization (validation on left, testing on right).

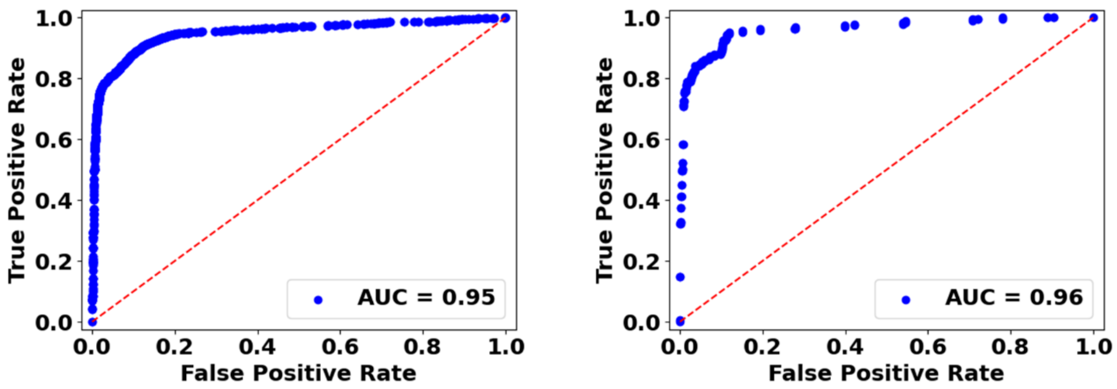


**Figure S21.** Receiver operating characteristic curve for extracellular localization (validation on left, testing on right).

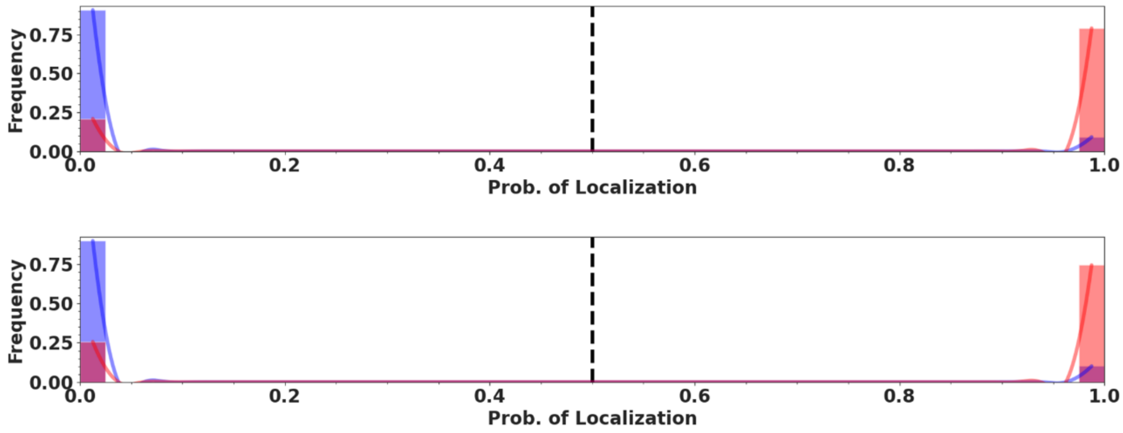


**Figure S22.** Classification histograms for nuclear localization (validation on top, testing on bottom).

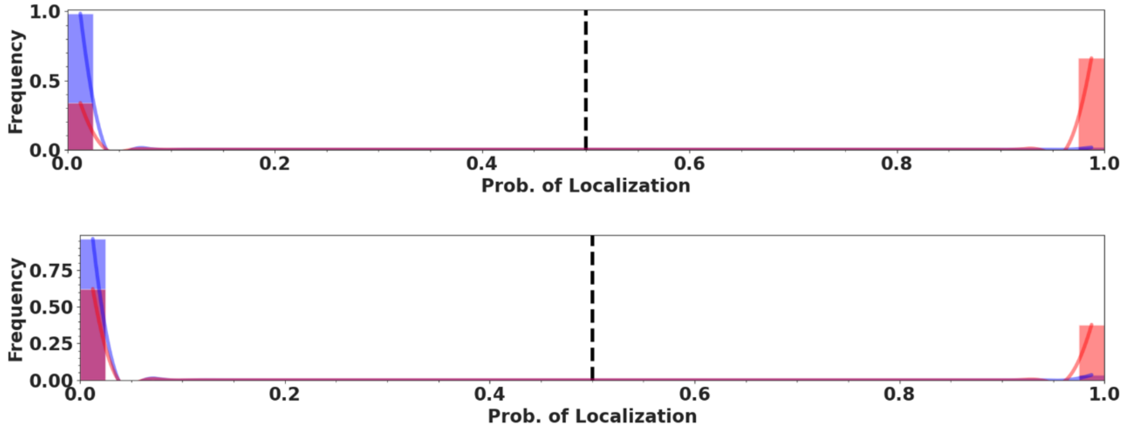


**Figure S23.** Classification histograms for mitochondria localization (validation on top, testing on bottom).

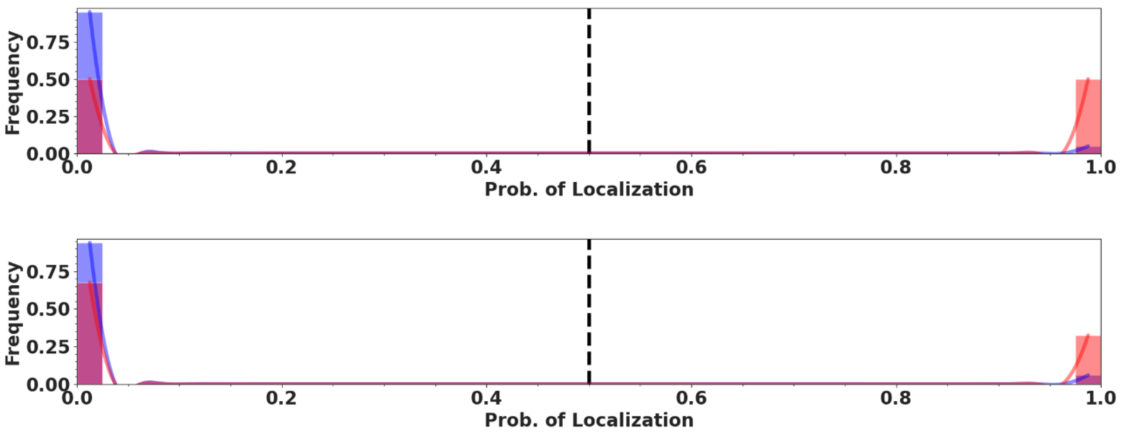


**Figure S24.** Classification histograms for endoplasmic reticulum localization (validation on top, testing on bottom).

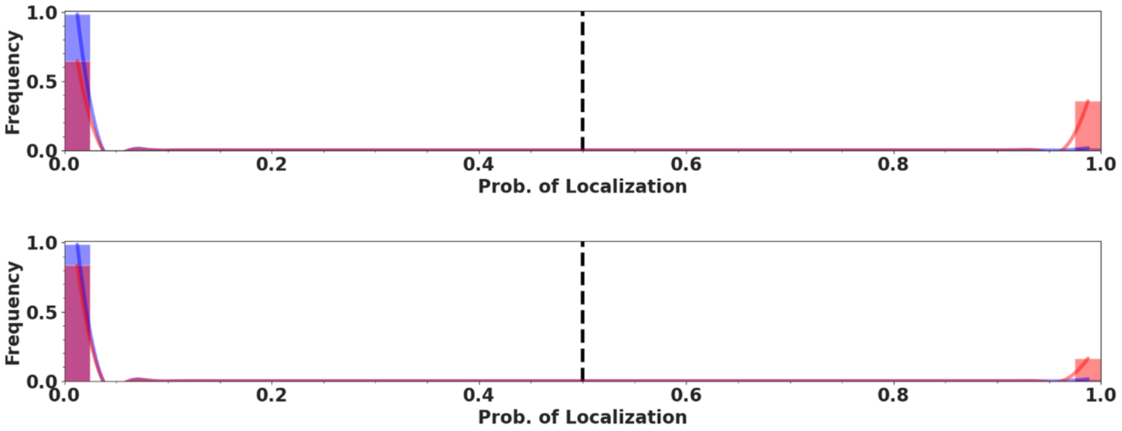


**Figure S25.** Classification histograms for golgi apparatus localization (validation on top, testing on bottom).

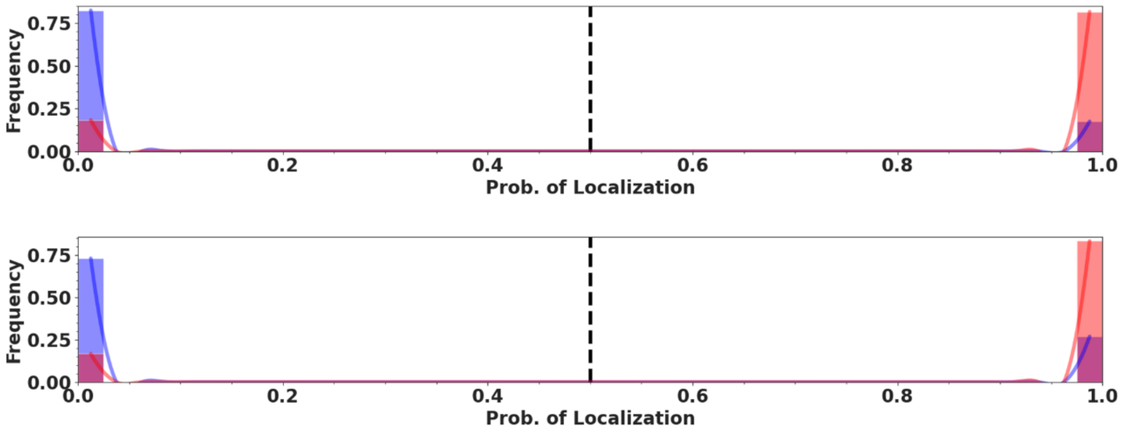


**Figure S26.** Classification histograms for cytoplasm localization (validation on top, testing on bottom).

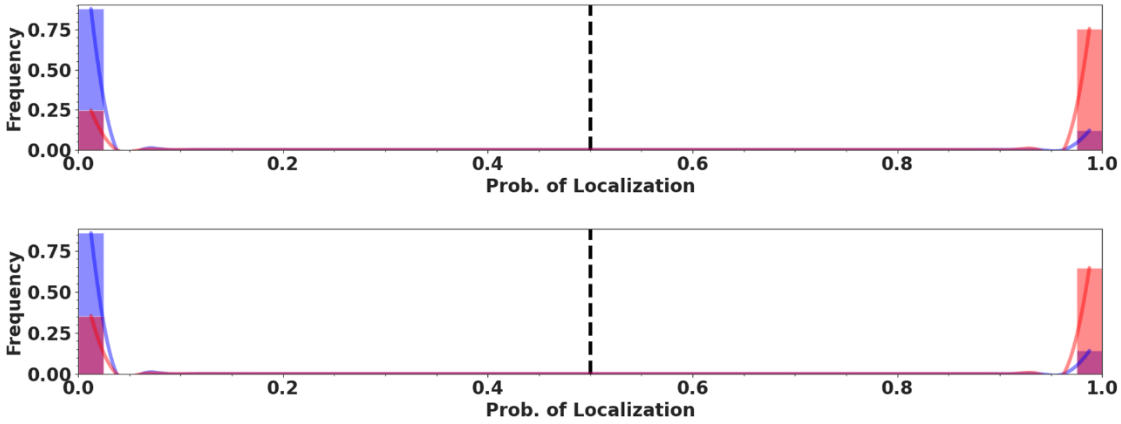


**Figure S27.** Classification histograms for cell membrane localization (validation on top, testing on bottom).

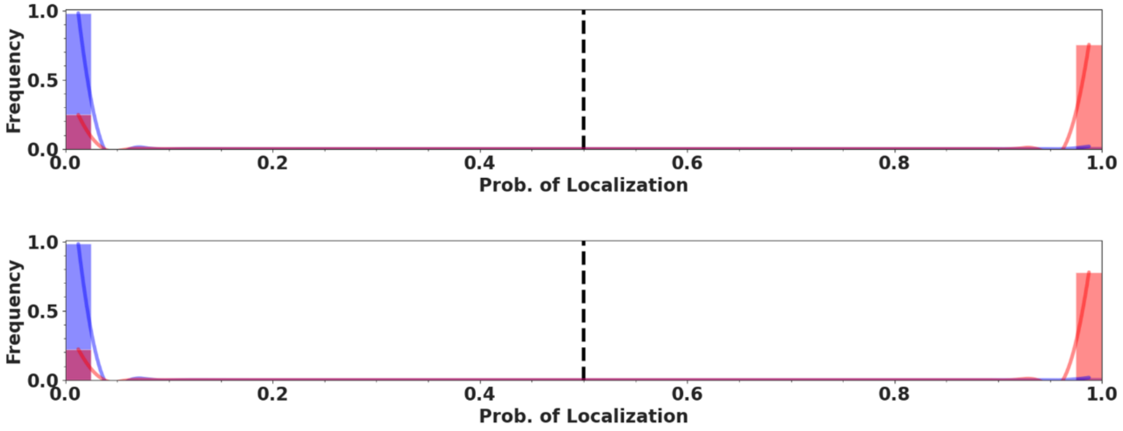


**Figure S28.** Classification histograms for extracellular localization (validation on top, testing on bottom).

**Table 1.** Table of testing F1 % change in Triplet models over ProtBERT model controls.

| Label | % F1 Increase |
|---|---|
| Nucleus | 3.90 |
| Mitochondria | 15.15 |
| Endoplasmic Reticulum | -8.33 |
| Golgi Apparatus | 71.43 |
| Cytoplasm | 0.10 |
| Cell Membrane | 1.79 |
| Extracellular Region | 15.71 |
